# DNA damage dependent hypomethylation regulates the pro-angiogenic LncRNA MEG9

**DOI:** 10.1101/442699

**Authors:** Cristina Espinosa-Diez, RaeAnna Wilson, Rishima Mukherjee, Marlee Feltham, Clayton Hudson, Rebecca Ruhl, Sudarshan Anand

## Abstract

Changes in gene expression are key for the cells to adapt and response to intrinsic and extrinsic stimulus. It has been shown that genotoxic stress induces global hypomethylation as a result of decreased expression of DNA methyl transferases (DNMT). We hypothesized that DNA damage suppresses long non-coding RNA expression in the vasculature via DNA methylation leading to more robust DNA repair/survival or cellular senescence/death cell fate decisions. We show here that ionizing radiation reduces the expression of DNMTs in the vascular endothelium and this leads to increased expression of the anti-apoptotic lncRNA MEG9. MEG9 is a lncRNA from the DLK1-DIO3 ncRNA cluster. Loss-of-function studies using RNA gapmers indicate that MEG9 protects endothelial cells from DNA damage induced cell death. Consistent with this phenotype, knockdown of MEG9 decreases growth factor dependent angiogenesis in a 3D fibrin gel angiogenesis assay. Mechanistically, we observed that MEG9 knockdown decreased the expression of cell survival genes including survivin and induced the expression of pro-apoptotic genes such as Bad/Bax. Taken together, our findings illustrate how DNA methylation at selective lncRNA loci can regulate their expression and drive endothelial cell fate decisions.

## Introduction

The cardiovascular system is highly sensitive to stress and one of the fastest to answer accordingly. Depending on how severe the damage is, endothelial cells (ECs) make quick decisions among cell death, cell cycle arrest or survival. We have demonstrated how the expression of specific group of miRs can influence endothelial fate [1, 2] in the context of DNA damage responses. However, the role of long non-coding RNAs (lncRNAs) in the endothelial stress responses is unclear.

LncRNAs are RNAs longer than 200 nt that regulate gene expression through epigenetic, transcriptional and post-transcriptional mechanisms but also interacting with functional proteins [3, 4]. Since lncRNAs are still a new class of regulatory molecules, extensive profiling of lncRNA expression across cell types, organs, or in response to extracellular stressors has not been done. Additionally, the potential role of lncRNAs in angiogenesis and their potential utility as targets for the development of therapeutics remains to be established [5, 6]. Consequently, we undertook a selective lncRNA screening approach to identify the differentially expressed lncRNAs across a number of organ systems and in response to angiogenic growth factors and several defined stressors.

We found that the LncRNA MEG9 is significantly upregulated after genotoxic stress in ECs. This lncRNA resides in the non-coding RNA cluster DLK1-DIO3, an imprinted genomic region that encodes not only lncRNAs but microRNAs [7, 8]. Although, several publications have described how DNA damaging agents and stressors modify methylation in this genomic region [9], it was not clear whether or not MEG9 was directly regulated by DNMT1. In this manuscript, we describd how methylation alters MEG9 expression and how MEG9 expression affects endothelial cell fates *in vitro*.

## Materials & Methods

### Cell Culture

HUVECs and HMVECs (Lonza) were cultured in EGM-2 media (Lonza) supplemented with 10% Fetal Calf Serum (Hyclone). HUVECs were treated with Cisplatin (5uM), Etoposide (10uM), Hydrogen Peroxide (500uM), Hydroxyurea (5uM), VEGF (50ng/mL), β-FGF (100ng/mL) and 10Gy γ-radiation from a cesium source. Cells were incubated at 37°C with 5% CO_2_ for 6 hours. VEGF was purchased from PeproTech, Inc. FGF was purchased from Fisher Scientific.

### Transfection

Cell were transfected at 50–60% confluence using standard forward transfection protocols using RNAimax reagent (Life Technologies) for siRNAs or gapmers. Typically, 50 nM RNA were used for transfections.

### RNA extraction and RT-PCR

Total RNA was isolated using a miRVana microRNA isolation kit (Ambion). Reverse transcription was performed using TaqMan™ Advanced cDNA Synthesis Kit (Life Tech) according to the manufacturer’s instructions. RT-PCR was performed using multiplexed TaqMan primers (Applied Biosystems). The relative quantification of gene expression was determined using the 2^-ΔΔCt^ method [10]. Using this method, we obtained the fold changes in gene expression normalized to an internal control gene, GAPDH or U6 snRNA, respectively.

### LncRNA Screen

RNA was diluted to 400ng/uL and cDNA synthesized in duplicate using the Systems Biology LncRNA profiler kit. The resulting cDNA was used in a SYBR-Green based qPCR to analyze the expression of 90 lncRNAs and 6 normalization genes (in duplicate). The data was analyzed, and Ct-values internally normalized to RNU43 to generate ΔCt-values, then treatment conditions were normalized to untreated (IgG treated) cells to generate ΔΔCt-values. Gene expression levels were converted to base-2 fold-change (2^-ΔΔCt^) [10], plotted and compared across conditions.

### Radiation of Cells

Cells were irradiation on a Shepherd□137cesium irradiator at a rate of ~166 cGy/min.

### Caspase-3/7 Activity & Cell Viability

Cells were transfected with Exiqon GapMers. After 48 hours, Caspase-3/7 activity and cell viability were measured using Promega Caspase-3/7 Glo and Cell Titer Glo 2.0 according to manufacturer’s protocol.

### BrdU Cell Proliferation Assay

Cells were transfected with Exiqon GapMers. After 24-hours BrdU was added to the media. The next day, cell proliferation was evaluated using the BrdU Cell Proliferation kit from EMD-Millipore. Colorimetric analysis was done with single channel 450nm intensity. Blank wells with either media alone or cells without BrDU were used for background correction.

### Western blot and densitometric analysis

After treatment, cells were washed in phosphate-buffered saline (PBS) and lysed in RIPA buffer (Sigma) supplemented with Complete Protease inhibitor cocktail (ROCHE) and Phosphatase inhibitors cocktail 2 and 3 (Sigma). Lysed cells were harvested by scraping, and proteins were analyzed by Western blot. Equivalent amounts of protein were loaded on a 4–12% gradient SDS-polyacrylamide gel (BioRAD) and transferred for 30 min in a TransBlot turbo (BioRAD) onto Nitrocellulose membranes. Membranes were blocked in 5% milk or 3% BSA and incubated with antibodies as indicated: DNMT1 (Cell Signaling, D63A6, 1:1000), DNMT3a (Novus, NB120-13888SS, 1:1000). GAPDH (Sigma, A5316, 1:10,000 1□h RT) was used as a housekeeping control for the total levels of protein loaded. Membranes were washed in TBST and incubated with secondary HRP antibodies from Cell Signaling Biosciences. HRP used were anti-mouse 7076P2 (1:5,000) and anti-rabbit 7074P2(1:5,000). Blots were scanned on the Licor Odyssey scanner according to manufacturer’s instructions. For protein array experiments, the human apoptosis array (R&D Biosystems ARY009) was used according to manufacturer’s instructions.

### 3-D Angiogenic Sprouting Assay

Early passage HUVECs were coated on Cytodex-3 beads (GE Healthcare) at a density of 10 million cells/40 μl beads and incubated in suspension for 3-4 hours with gentle mixing every hour. They were plated on TC treated 6 well dishes overnight and resuspended in a 2mg/ml fibrin gel with 200,000 human vascular smooth muscle cells. The gel was allowed to polymerize, and complete EGM-2 media was added. Sprouts were visualized from days 3-4 via confocal imaging after overnight incubation with FITC labeled Ulex europaeus lectin (Vector labs). Immunofluorescence imaging was performed on a Yokogawa CSU-W1 spinning disk confocal microscope with 20x, 0.45 Plan Fluor objective (Nikon).

### Statistics

All statistical analysis was performed using Excel (Microsoft) or Prism (GraphPad). Two-tailed Student’s *t*-test or Mann–Whitney *U*-test was used to calculate statistical significance. Data that was not normally distributed as assessed by Shapiro-wilk test (Excel, Real statistics add-in) was evaluated using *U*-test. Variance was similar between treatment groups. A *P* value<0.05 was considered to be significant.

## Results

### Pro-angiogenic growth factors alter LncRNA expression

Several publications have previously described the increasing role of lncRNAs in physiological and pathological angiogenesis [11-13]. Vascular endothelial growth factor (VEGF) and fibroblast growth factor β (β-FGF) are key regulators in the angiogenic process, during development or in adulthood in response to damage or a wound. Therefore, we screened for lncRNAs induced by both angiogenic growth factors in human umbilical vein endothelial cells (HUVECs). Among the 90 LncRNAs tested, we identified four lncRNA that were upregulated by VEGF and β-FGF. The first up-regulated lncRNA is EGO-A/B (EGOT), associated with malignant breast cancer and that inhibits the anti-viral response to hepatitis C virus [14]. In the same group, we identified H19-up-stream: a previously identified exosomal regulator of endothelial cells that promotes tube formation. WT1-AS, the anti-sense transcript of Wilm’s tumor associated gene 1 (WT1), negatively regulates WT1. WT1-AS down-regulation promotes lung metastasis in gastric cancers [15] (Fig 1A). Interestingly, only one lncRNA was down regulated by both VEGF and β-FGF, the maternally imprinted gene 9 (MEG9) (Fig 1B). MEG9, also known as Mirg in mouse, is a maternally imprinted gene expressed in embryonic tissue with some context specific expression in adult tissue [16, 17]. Aberrant repression of Meg9 and other maternally expressed lncRNAs from the DLK1-Dio3 imprinting cluster is present in most induced pluripotent stem cell (iPSC) lines and is thought to be responsible for the failure of iPSCs to form viable mice, with embryos dying mid-gestation [18].

**Figure 1:**
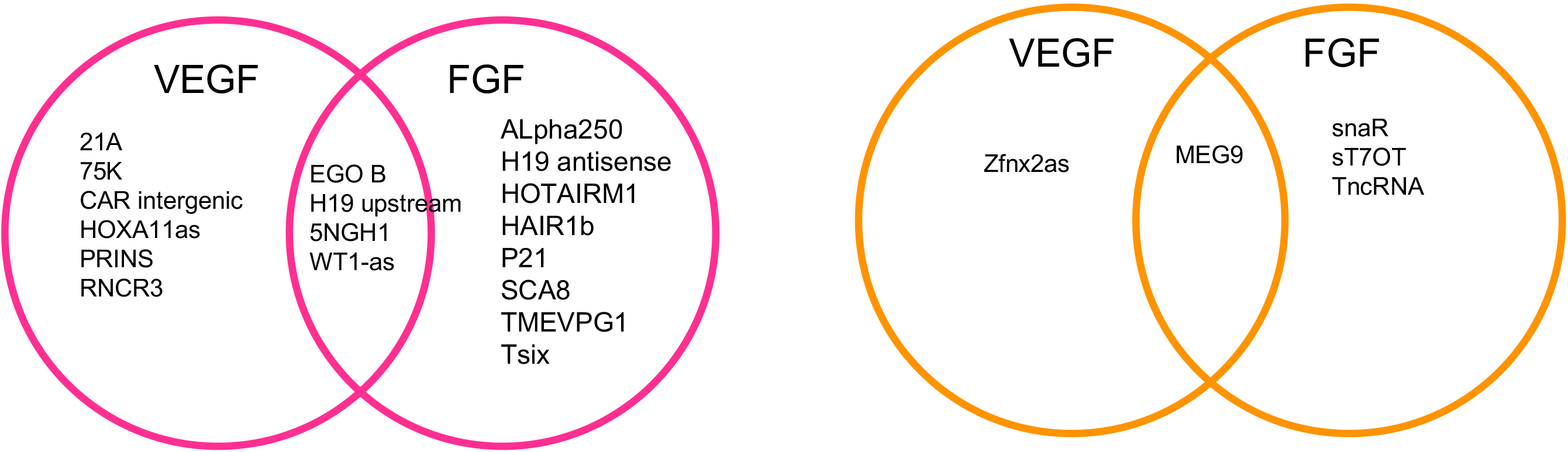
Venn Diagrams of lncRNAs upregulated (pink) and downregulated (orange) after VEGF and FGF treatment. HUVECs were plated at 70% confluency and starved overnight in endothelial basal medium (EBM2). After starvation, cells were treated with 50 ng/ml of VEGF or 100 ng/ml of bFGF for 6 hours. LncRNA expression was profiled using LncRNA arrays. HUVECs in EBM2 were used as control.

### DNA Damaging Agents and Replication Stressors Regulate the Same Sets of lncRNAs

Cell stressors often alter the expression profiles of both coding and non-coding RNAs in order to respond to the stress [19, 20]. Depending if the damage is severe or moderate, the cells will respond differently. To identify how ECs respond to different stressors (DNA damage, ER stress, oxidative stress), we examined the expression profiles of HUVECs exposed to the oxidative stressor hydrogen peroxide, and to double-strand break-inducing replication fork stressors cisplatin, etoposide, hydroxyurea and a 10 Gy dose of Ionizing Radiation (IR). We found that 5 lncRNAs were up regulated in common in all treatment conditions (Fig 2). WT1-AS was more than 8-fold up regulated in all treatments. This is an interesting finding, as WT1-AS was also up regulated with VEGF and β-FGF treatment. Other lncRNAs up regulated by all four treatments were Tsix, NDM29 (neuroblastoma differentiation marker 29), snaR and Y-RNA-1. NDM29 has previously been found to be up regulated by cytotoxic substances and has been found to inhibit cancer proliferation by promoting terminal differentiation in Neuroblastoma [21].

**Figure 2:**
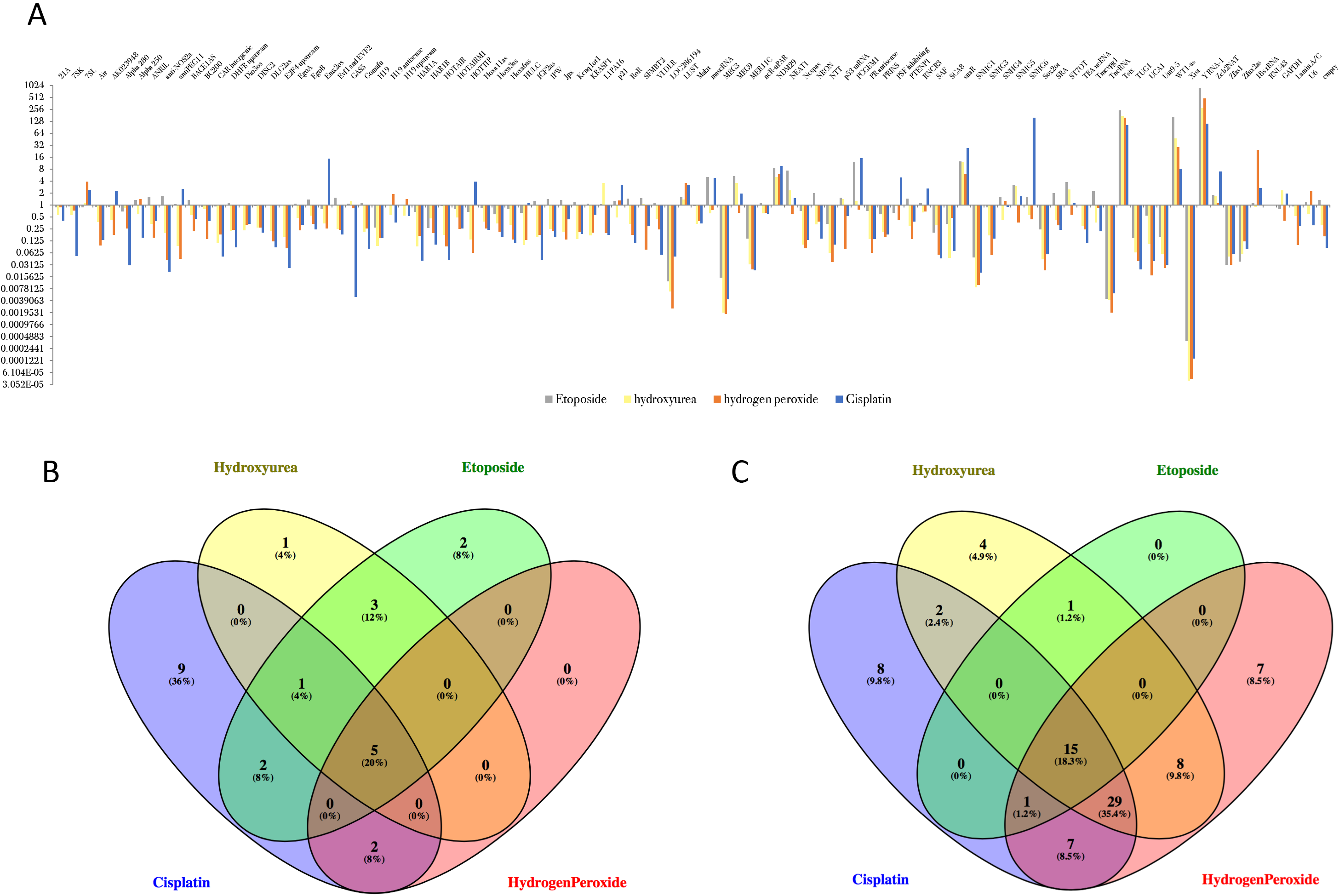
LncRNA changes in endothelial cells after DNA damage. A. Chart of fold-enrichment of lncRNAs in indicated treatment group Cisplatin (5uM), Etoposide (10uM), Hydrogen Peroxide (500uM), Hydroxyurea (5uM) relative to untreated cells. B. Venn diagram of enriched lncRNAs relative to untreated control. C. Venn diagram of depleted lncRNAs relative to untreated control.

Furthermore, MEG9 was upregulated with all the treatment but hydrogen peroxide and we observed a similar increase with ionizing radiation, in contrast to our previous results with VEGF and bFGF. We decided to focus on the role of MEG9 in ECs and the regulation of its expression. Interestingly, the chromosomal region that MEG9 resides in, the DLK1-Dio3 cluster (Fig 3A) has been shown to be regulated by DNA methylation [22]. The mouse and human loci are both regulated by a 5’ intergenic differentially methylated region (IG-DMR). LncRNAs are expressed from the non-methylated maternal allele. In contrast, the paternal chromosome is methylated preventing the expression of the ncRNAs from this cluster [23].

**Figure 3:**
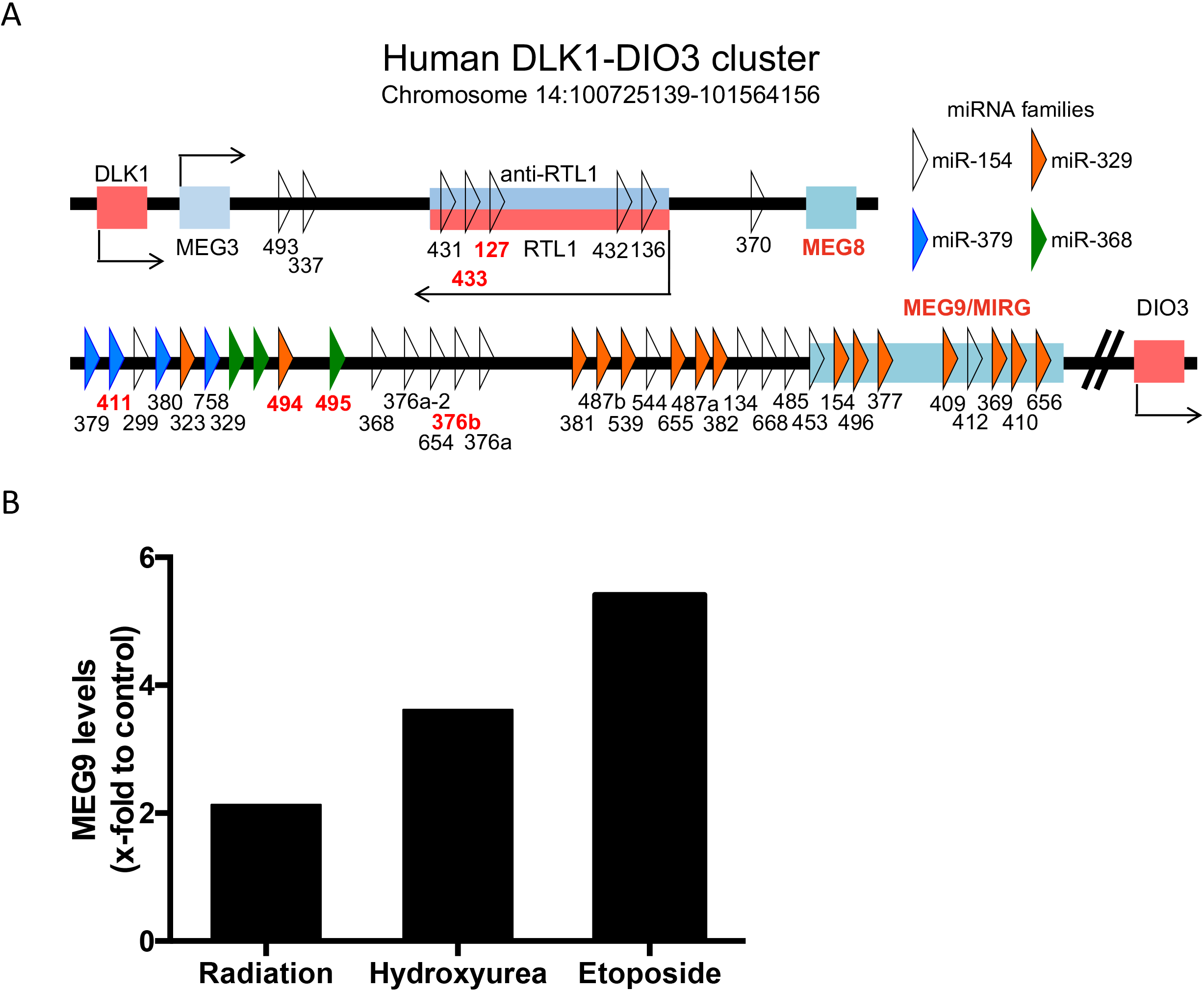
Schematic representation of the 1-Mb imprinted cluster on human chromosome 14. Representation of the location of lncRNAs and miRNAs in the locus. Paternal (blue) and maternal (pink). microRNAs and LncRNAs in red have been associated with DNA damage in previous studies. (Adapted from Kircher et al 2008). B. Meg 9 expression levels. HUVECs were treated with Radiation (10 Gy), hydroxyurea (5 uM) or etoposide (10 uM) for 6h. RNA was analyzed by qPCR using GAPDH as a housekeeping control.

### Genotoxic stress triggers changes in DNA methylation affecting Meg9 expression

Demethylation of the DLK1-DIO cluster has been previously described in human patients exposed to stressors. Smokers that have developed lung cancer showed differences in methylation in lncRNAs from this cluster such as MEG3, MEG8 and MEG9 [9]. We hypothesized that the significant upregulation of MEG9 we observed after genotoxic stress would be dependent on decreased DNA methylation. We first established that IR decreases expression of DNMT1 and DNMT3a (Fig 4A-B), along with increased expression of the DNA demethylases TET1 and TET3 (Fig 4C) in a dose dependent manner. Interestingly, knockdown of DNMT1 using gapmers (Sup Fig 1) mimics radiation by driving expression of MEG9 (Fig 4D). Together this data suggests that MEG9 is epigenetically regulated upon stress through demethylation.

**Figure 4:**
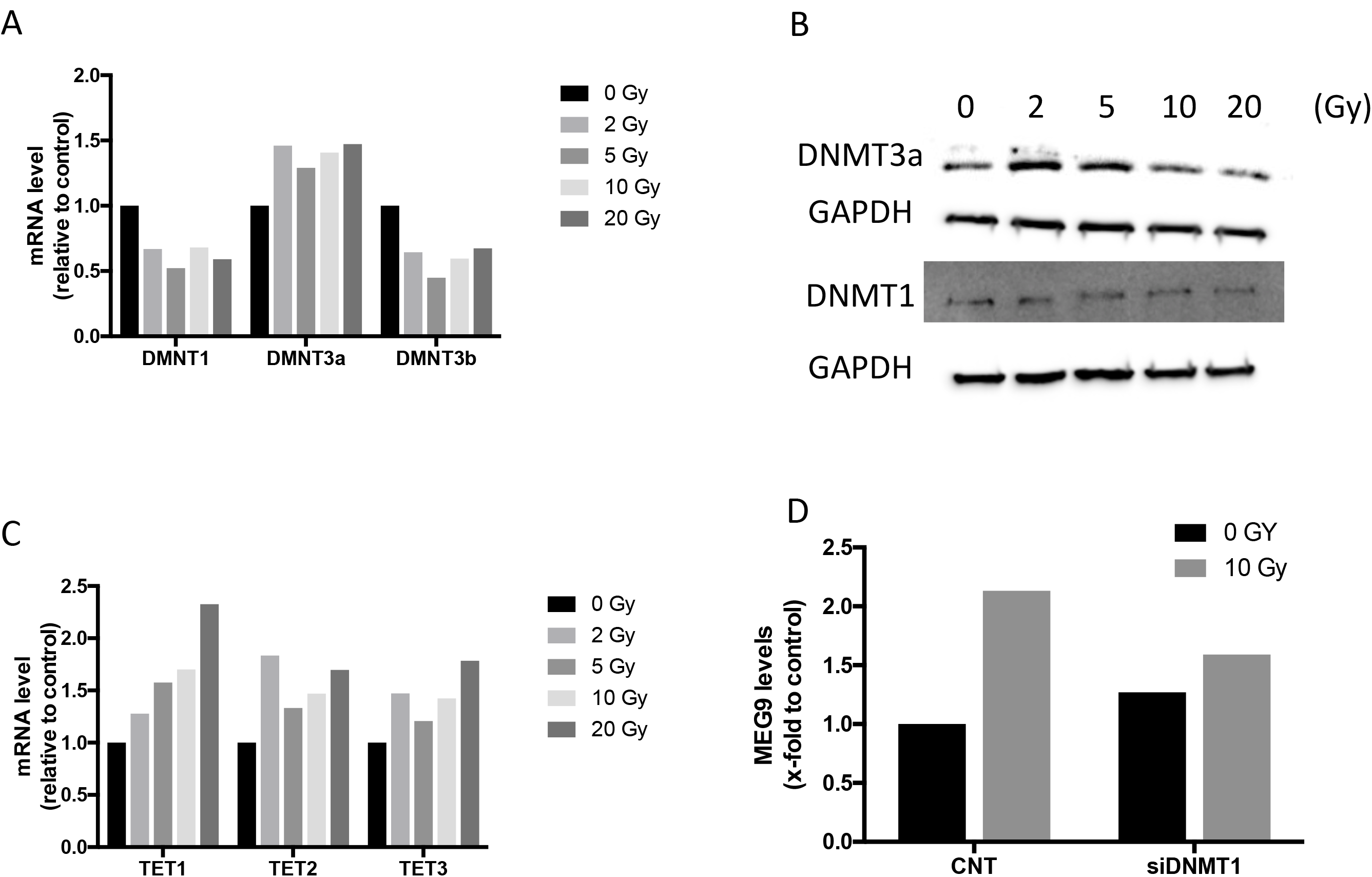
DNA damage regulates DNA methylation. A. RNA expression of DNMT1, DNMT3a and DNMT3b. Bar graphs represent mRNA levels relative to housekeeping GAPDH. B. Representative western blot of DNMT1 and DNMT3a in HUVECs treated with different doses of Ionizing Radiation. C. mRNA expression of TET family demethylases. HUVECs were irradiated with different doses of radiation (0, 2, 5, 10 and 20 Gy) and incubated for 24h. D. Meg 9 expression levels. HUVECs were transfected with DNMT1 siRNA for 24 hours and irradiated with 10 Gy. After 24 h cells were harvested and RNA was analyzed by qPCR.

### Meg9 loss of function leads to increase apoptosis and decreases endothelial cell proliferation

We have previously described how acute genotoxic stress stimulus lead to apoptosis and high doses of IR lead to EC cell death [2]. As MEG9 function in adult EC has not been described in depth, we tested if it has a role in triggering cell death. We decreased MEG9 levels in HUVECs using a validated siRNA gapmer (Sup Fig 2) and we assayed cell death and cell proliferation using standard assays (Fig 5A-5B) and BrdU staining (Fig 5C). Inhibition of MEG9 induces a decrease in cell proliferation and viability, that correlates with an increase in cell death. To explore the mechanisms behind these effects, we analyzed the protein levels of key apoptotic markers using an apoptosis protein array. Compare to the control treated ECs, MEG9 inhibition increase the levels of the pro-apoptotic proteins Bad and Bax, 75 and 50-fold respectively and reduces the levels of cytoprotective proteins such as the antioxidants HO-1 and HO-2 (Fig 6).

**Figure 5:**
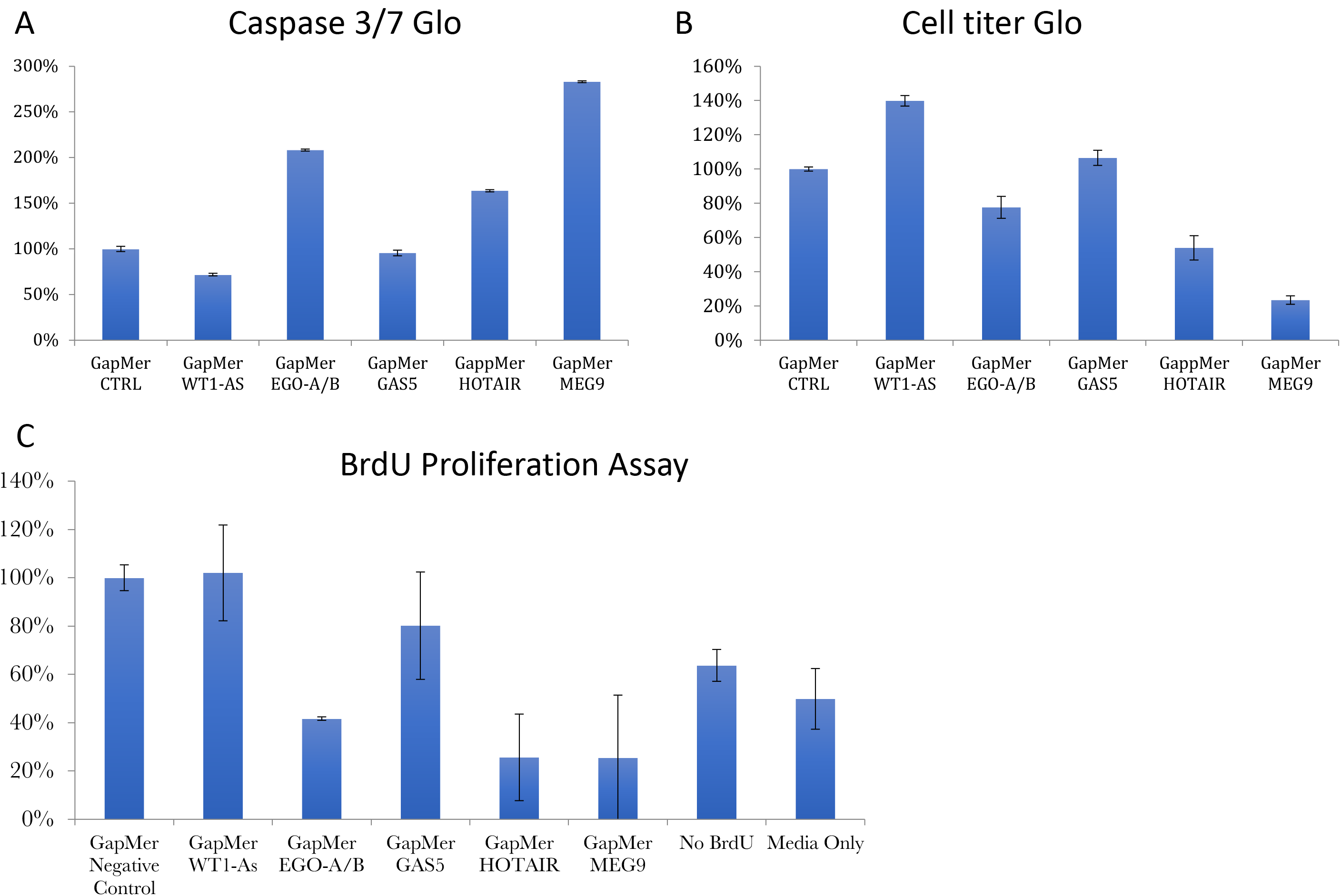
Functional role of lncRNAs in endothelium. A. Caspase 3/7 glo assay. HUVECs were transfected with the indicated lncRNA gapmers and after 48h cell death was assayed using Casp3/7 glo. Bar graphs depict mean+SD. B. % Proliferation in HUVECs transfected for 48 h with the indicated Gapmers against LncRNAs. Bar graphs depict mean+ SD. C. BrdU staining in HUVECs after 48 h transfection with LncRNA gapmers. BrdU staining was evaluated by ELISA kit. Bar graph means indicate average of three technical replicates.

**Figure 6:**
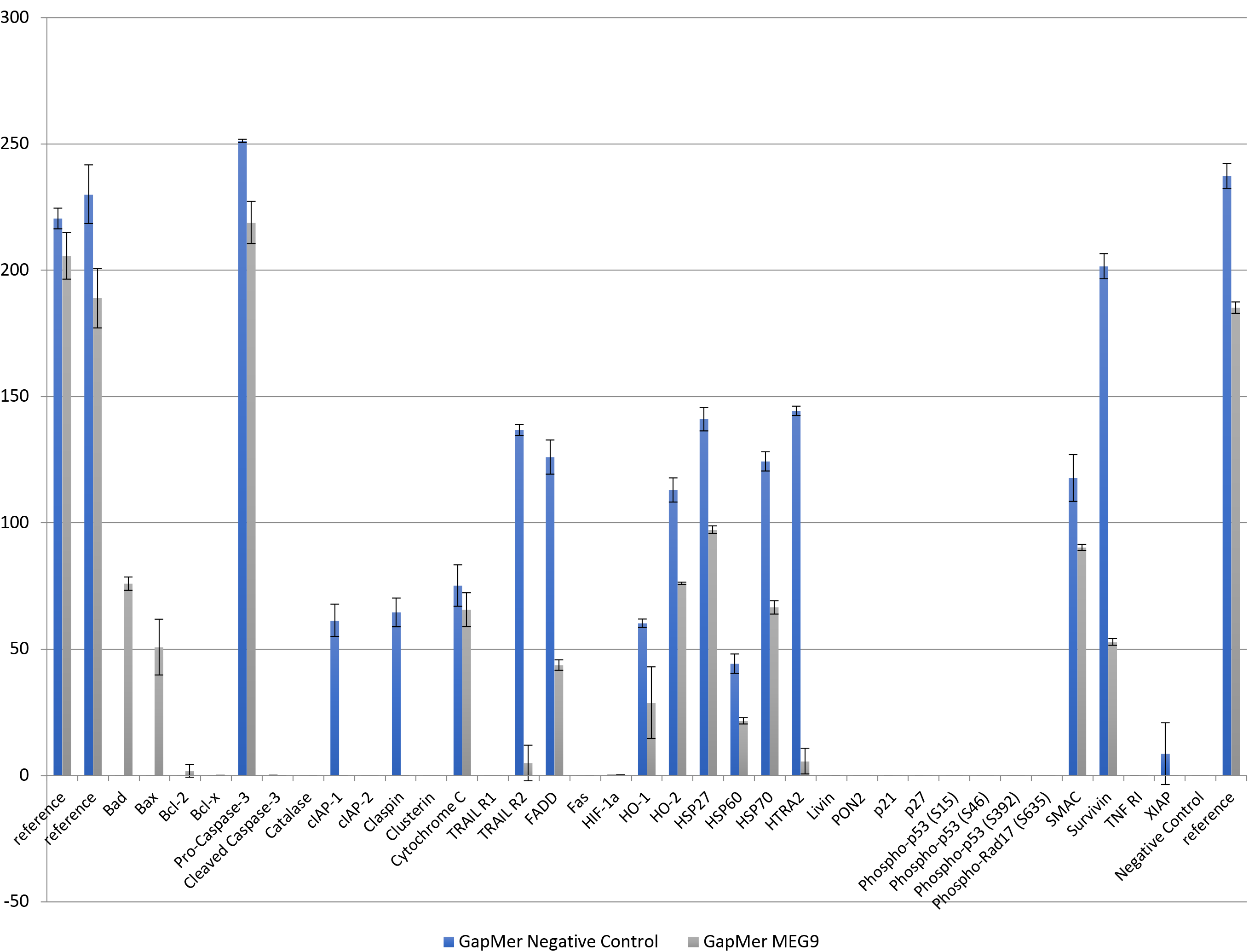
MEG9 inhibition induces expression of proapoptotic proteins. Representative quantitation of an apoptotic protein array. HUVECs were transfected with MEG9 or Control Gapmer for 48h. Protein array membranes were incubated with 1 ug of protein and develop using HRP. Bar graphs show mean+ SD of two independent replicates.

### Inhibition of MEG9 leads to decrease of sprouting angiogenesis in vitro

Angiogenesis in response to GF is a reliable way to measure EC health and fitness. Using a cytodex bead based 3D sprouting angiogenesis assay we evaluated the role of MEG9 in sprouting angiogenesis. Consistent with our other readouts, MEG9 inhibition significantly decreased the area of sprouts on day 5 in our *in vitro* model, suggesting that MEG9 would have a pro-angiogenic role (Fig 6). Although, the expression of MEG9 decreases with VEGF or FGF treatment suggesting a negative role, we hypothesize that MEG9 is a pro-angiogenic lncRNA based on the cell based 2D and 3D assays. It is also possible that MEG9 is acting in an inhibitory feedback loop during vascular damage and the direct expression levels are not reflective of its biological function (Fig 7).

**Figure 7:**
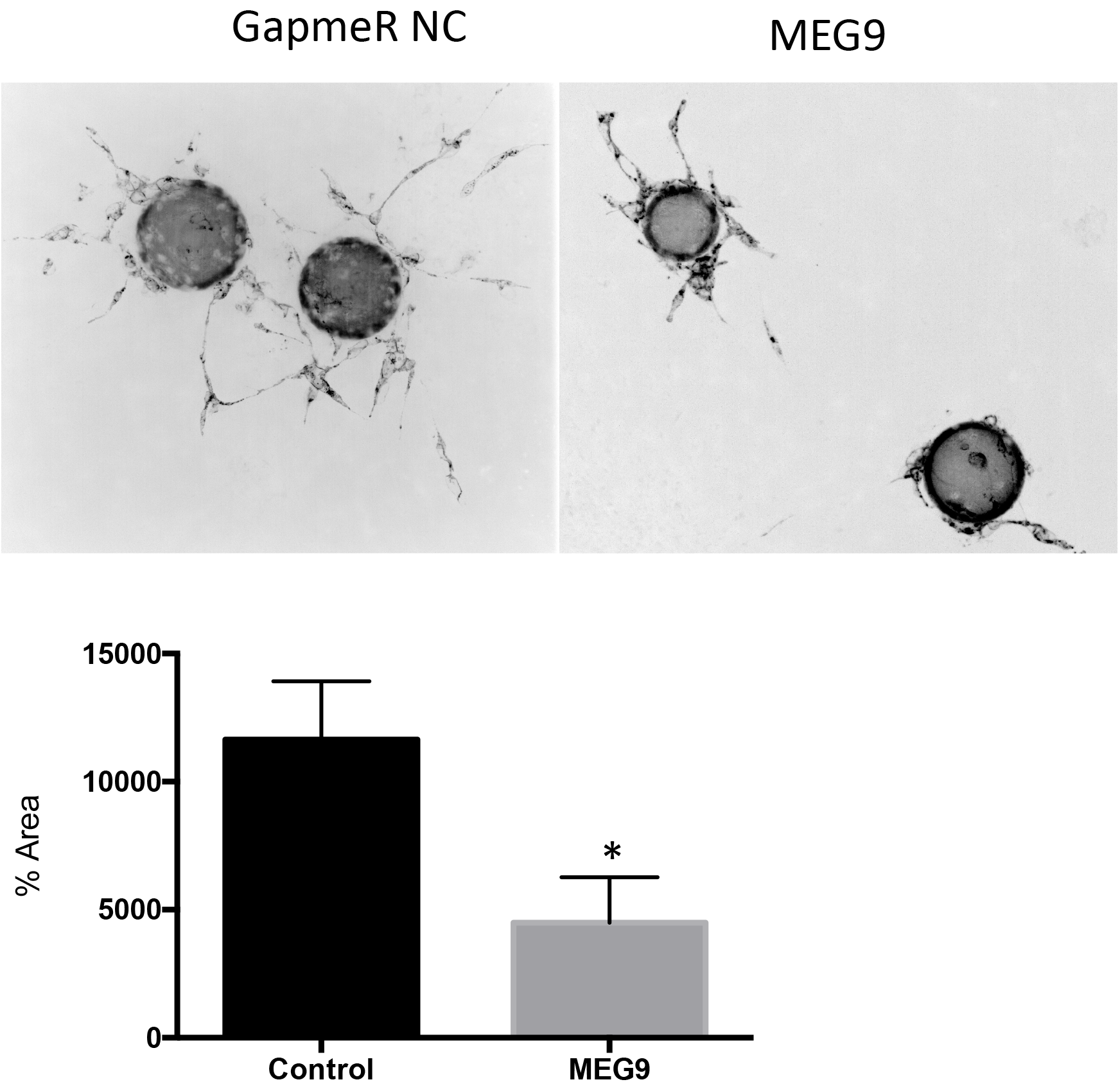
MEG9 inhibition reduces sprouting angiogenesis. Fibrin bead angiogenesis assay. HUVECs were transfected with the indicated siRNA gapmers and assessed for their sprouting angiogenesis potential. The images show representative lectin-stained beads (green) for each condition. Bars depict mean + SEM of lectin area analyzed across at least 25 beads/group. *P < 0.05; two-tailed Student’s T-test

**Figure 8:**
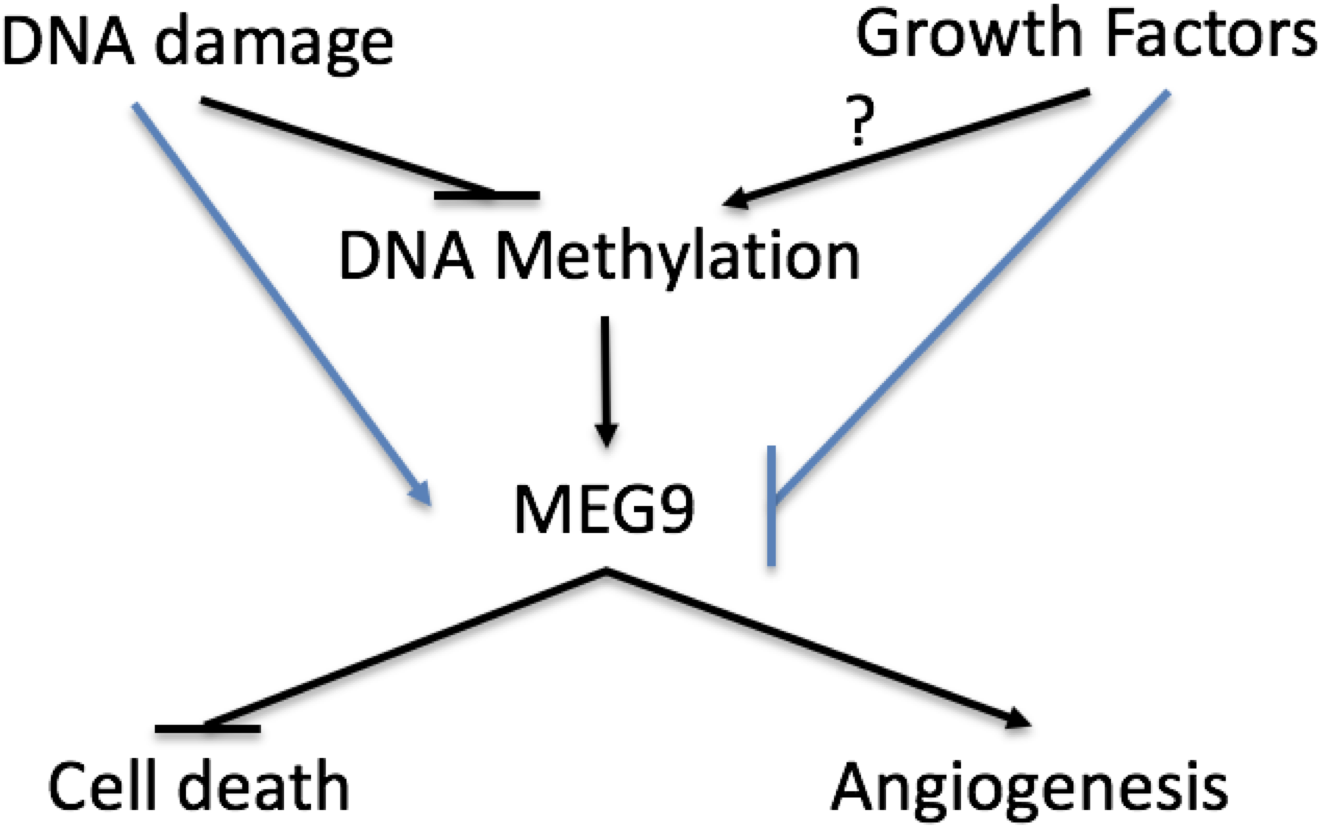
Proposed model. Protective role of MEG9 in endothelium. MEG9 is upregulated after DNA damage and genotoxic stress but downregulated in response to Growth Factors such as FGF and VEGF. This leads to a pro-survival and pro-angiogenic role.

## Discussion

Our current investigation suggests that DNA damage induce hypomethylation is able to regulate lncRNAs from the DLK1-DIO3 cluster, such as MEG9 through inhibition of DNMT expression. In the absence of TET1, MEG9 induction after DNA damage is blocked, indicating that active role of the DNA demethylases Ten-eleven translocation enzymes (TET) is needed. This is not unexpected either, as TET enzyme expression increased after DNA damage.

Kamesmaran et al confirmed how susceptible the DLK1-DIO3 cluster is to methylation by targeting the specific promoter of lncRNA MEG3 in islets from type 2 Diabetes patients [24]. We have observed in our profiles that MEG9 and MEG3 respond in opposite directions to DNA damage and growth factor responses. While MEG3’s role in vascular pathology has been broadly described, the role and function of MEG9 is still unknown. Interestingly, Boon et al have described MEG3 was significantly upregulated in senescent HUVECs and in human cardiac atria from aged patients’ samples. Loss of function studies with MEG3-LNA increased sprouting angiogenesis *in vitro* and perfusion *in vivo* [25] and ([26]. In myocardial infarction, MEG3 was also upregulated and a trigger for cell death and cardiomyocyte apoptosis [27].

These studies together with our results point out two lncRNAs encoded in the same megacluster are epigenetically regulated through DNA methylases and with opposite and complementary modulators. Whether or not MEG9 and MEG3 balance each other inducing and regulating opposite pathways in angiogenesis needs further investigation. At this point, we can conclude is MEG9 inhibition *in vitro* leads to endothelial cell death and decreases sprouting angiogenesis. Loss of function of MEG9 inhibits key proapoptotic proteins such as Bad, Bax and Bcl2 [28, 29] while increasing cell death. We propose that further investigation into the regulation of MEG9 and its function in vascular cells will provide insight into how lncRNAs in the DLK3-Dio cluster can direct different cell fate programs.

---

**Supplementary Figure 1:**
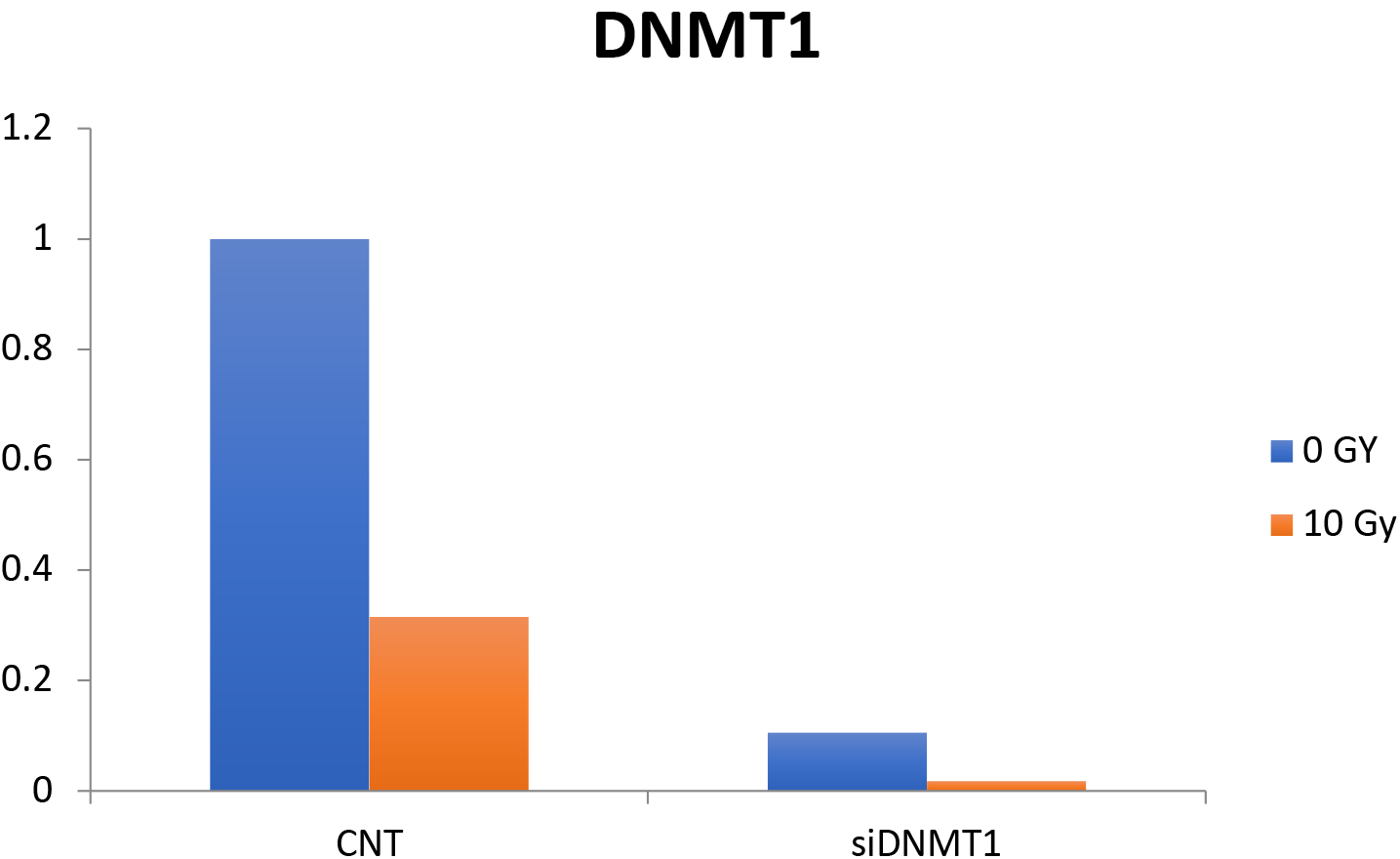
Validation of DNMT1 gapmer. HUVECs were transfected with Gapmer control or DNMT1 for 24 h and then treated with 10 Gy for other 24 h Expression levels of DNMT1 were analyzed by PCR using GAPDH as housekeeping control.

**Supplementary Figure 2:**
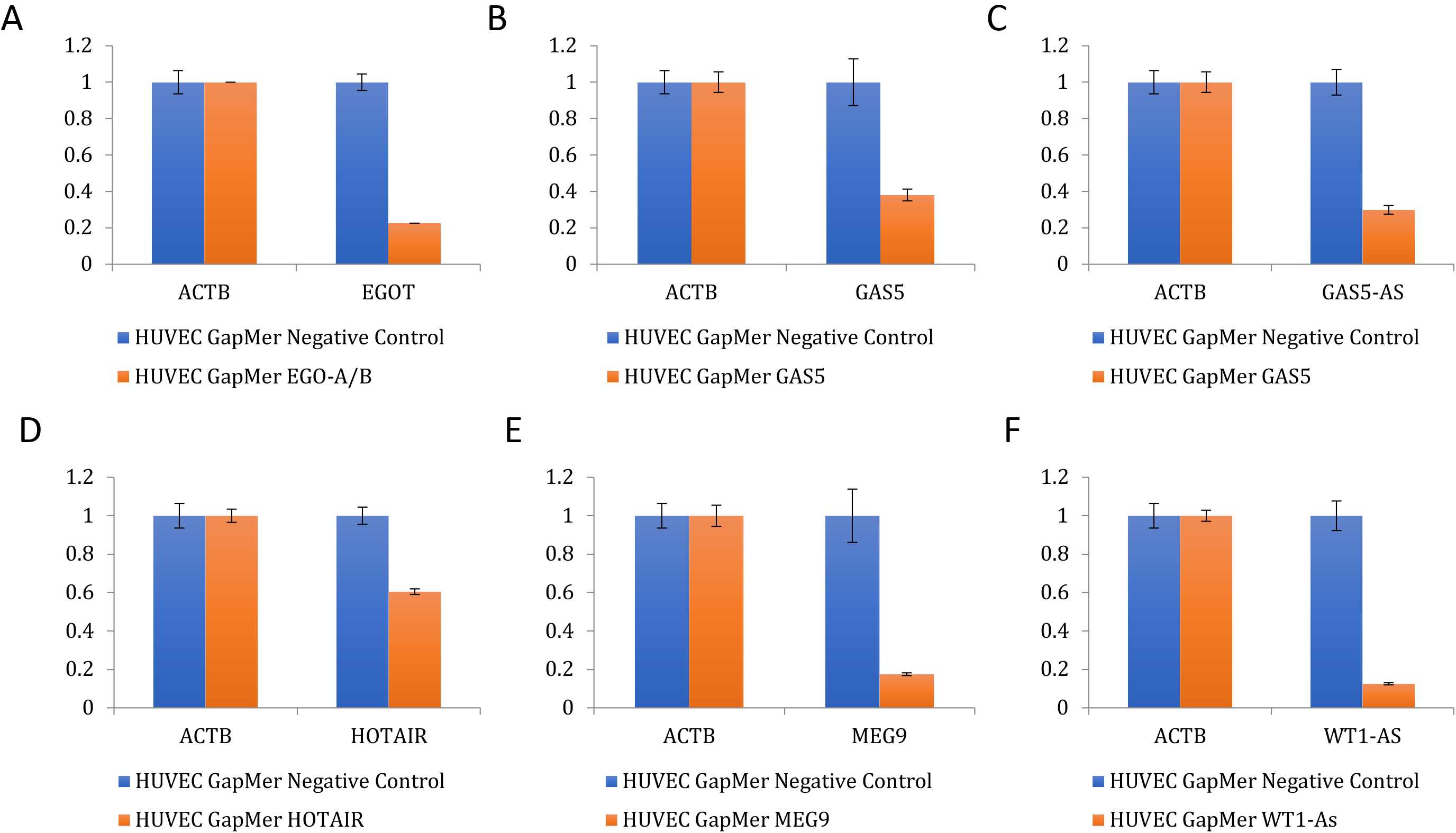
Validation of LncRNA Gapmers. HUVECs were transfected with A. EGOT, B. GAS5, C.GAS5-AS, D. HOTAIR, E. MEG9 and F. WT1-As for 24h. Expression levels of lncRNAs were analyzed by PCR using GAPDH as housekeeping control. Bar graphs depict mean+SD.

